# High-frequency temperature variability mirrors fixed differences in thermal limits of the massive coral *Porites lobata* (Dana, 1846)

**DOI:** 10.1101/367763

**Authors:** DJ Barshis, C Birkeland, RJ Toonen, RD Gates, JH Stillman

## Abstract

Spatial heterogeneity in environmental characteristics can drive adaptive differentiation when contrasting environments exert divergent selection pressures. This environmental and genetic heterogeneity can substantially influence population and community resilience to disturbance events. Here, we investigated corals from the highly variable back reef habitats of Ofu Island in American Samoa that thrive in thermal conditions known to elicit widespread bleaching and mortality elsewhere. To investigate the hypothesis that thermal variability is the driving force shaping previously observed differences in coral tolerance limits in Ofu, specimens of the common Indo-Pacific coral *Porites lobata* (Dana, 1846) from locations with differing levels of thermal variability were acclimated to low and high thermal variation in controlled common garden experimental aquaria. Overall, there was minimal effect of the acclimation exposure. Corals native to the site with the highest level of daily variability grew fastest, regardless of acclimation treatment. When exposed to lethal thermal stress, corals native to both variable sites contained elevated levels of heat shock proteins and maintained photosynthetic performance for 1–2 days longer than corals from the stable environment. Despite being separated by < 5 km, there was significant genetic differentiation among coral colonies (*F_ST_* = 0.206, p < 0.0001; nuclear ribosomal DNA), while *Symbiodinium* phylotypes were all ITS2-type C15. Our study demonstrates consistent signatures of adaptation in growth and stress resistance in corals from naturally thermally variable habitats, emphasizing that existing genetic diversity of corals is an important asset in strategies to protect and manage coral reef ecosystems in the face of global change.

**Summary Statement:** Corals native to highly variable habitats demonstrate greater thermal tolerance than corals from less variable habitats after 36-days of acclimation to thermally stable or variable common garden treatments.

## Introduction

Heterogeneous environments can drive adaptive diversification when contrasting environmental conditions exert strong divergent selection pressures and individual niches are not abundant enough to favor the evolution of overall plasticity (e.g., Dempster, 1955; Kawecki and Ebert, 2004; Levene, 1953; Ravigné et al., 2004). Local adaptation is expected to evolve in populations with limited connectivity, but if environmentally driven selection is strong enough, adaptive differentiation can still accumulate despite ongoing gene flow (Feder et al., 2012; Hoey and Pinsky, 2018). In the marine environment, reproduction via broadcast spawning and gamete mixing at the sea surface means dispersal potential (i.e., gene flow) among neighboring habitats can often be high, with larval neighborhoods sizes of many marine organisms estimated at > 10 km (e.g., Palumbi, 2004; Pinsky et al., 2017). Thus, for small-scale population differentiation to be driven by selection in the sea, certain genotypes must preferentially settle in optimal habitat-types, or sub-optimal settlers must have reduced fitness via strong post-settlement selection (Dempster, 1955; Levene, 1953; Ravigné et al., 2004).

Despite an established theoretical framework, the functional dynamics of adaptation and natural selection in most species remain unknown and these processes are particularly complex in reef building corals due to the symbiotic nature of the organism (Baird et al., 2007; Pandolfi et al., 2011). For example, adaptation of coral endosymbiotic algae, *Symbiodinium* spp., is known to confer varying degrees of thermal tolerance (Howells et al., 2012), and *Symbiodinium* diversity within individual host coral species can vary across thermal environments (D’angelo et al., 2015; Oliver and Palumbi, 2011). The specifics of how the genetic diversity of the coral host contributes to adaptation, however, is relatively unknown (Baird et al., 2009; Baird et al., 2007; Barshis et al., 2010; Dixon et al., 2015; Hoegh-Guldberg et al., 2007; but see Lundgren et al., 2013; Matz et al., 2018). Adaptation to environmental change, including climate shifts, has been demonstrated in other organisms (Hancock et al., 2011; Hoffmann and Sgro, 2011; Sanford and Kelly, 2011), and recent evidence for corals suggests that adaptive differences in coral thermal tolerance are heritable (Dixon et al., 2015; Kenkel et al., 2015; Meyer et al., 2009); lending credence to the idea of evolutionary rescue (sensu Bell and Gonzalez, 2009) of corals from climate impacts.

The back reef pools on Ofu Island, American Samoa, represent a natural laboratory for investigations of adaptation and acclimatization of corals to contrasting environments due to their high diurnal variation and small-scale heterogeneity in environmental characteristics (e.g., temperature, pH, flow, dissolved O2; Barshis et al., 2010; Craig et al., 2001; Smith et al., 2007). For example, the highly variable (HV) back reef pool of Ofu undergoes daily temperature fluctuations of up to 5.6 °C and reaches daily extremes of > 35 °C (mean daily range 1.59 °C ± 0.42 SD). In contrast, the adjacent less variable forereef has seasonal maximum daily temperature fluctuations of 1.8 °C (mean daily range is 0.6 ± 0.2 SD; Craig et al., 2001; Smith et al., 2008; unpublished data). Corals from among these thermal habitats have phenotypic differences consistent with local adaptation of thermal performance, including increased prevalence of heat-tolerant clade D *Symbiodinium* (e.g., Acropora spp., Pocillopora spp., Pavona spp.; Cunning et al., 2015; Oliver and Palumbi, 2009), constitutive turning-on of genes involved in cellular stress defense (Barshis et al., 2013), fixed and plastic responses following field transplantation (Palumbi et al., 2014; Smith et al., 2007; Smith et al., 2008), and small-scale (< 5 km) genetic differentiation of coral hosts (Barshis et al., 2010; Bay and Palumbi, 2014).

In the massive coral *Porites lobata* (Dana, 1846), host genotypes were subdivided across small spatial scales (< 5 km), while all *Symbiodinium* sequences matched ITS2 phylotype C15 (Barshis et al., 2010). The genetic differentiation of the host mirrored fixed differences in the cellular stress response (Barshis et al., 2010) and growth characteristics (Smith et al., 2007) suggestive of genetic adaptation to differences in the amount of diurnal environmental variability between back-reef pools; however, upper thermal limits were not tested in previous *P. lobata* studies. Here, we explore whether high-frequency thermal variability (defined here as diurnal or shorter variation *sensu* Safaie et al., 2018) is the environmental factor that differentiates growth and thermal tolerance of *P. lobata* colonies from contrasting habitats on Ofu. We used a common-garden laboratory acclimation experiment to test the hypothesis that corals from different thermal habitats have unique responses to daily thermal variation.

## Materials and Methods

### Study site, sample collection and transport

Corals were collected during May 2007 from three sites on Ofu and Olosega islands in the territory of American Samoa (14˚11’ S, 169˚40’ W). These islands host diverse communities of ∼85 shallow reef-building coral species, many of which are consistently exposed to atypically high seawater temperatures (Craig et al., 2001) and irradiances (Smith and Birkeland, 2007). Two back reef sites, a High Variability (HV) and Medium Variability (MV) pool, and one low-variability forereef site (forereef) were selected based on general differences in environmental characteristics (Craig et al., 2001; Piniak and Brown, 2009; Smith et al., 2007; Smith and Birkeland, 2007; Smith et al., 2008). Briefly, the HV pool is smaller, shallower, more thermally variable, and experiences higher water flow than the MV pool, while the forereef is relatively more stable than the HV and MV pools.

A pneumatic hole saw drill was used to remove n=30, 19 mm diameter cores from the upward facing surfaces of each of n=5 source colonies in each site (total n=150 cores). Source colonies were of similar size (1–1.5 m diameter) and at least 5 m apart to minimize potential for sampling the same clone (i.e., genet). Cores were affixed to nylon bolts with Z-Spar, Splash Zone marine epoxy (Carboline Company, St Louis, MO) and placed in the MV pool for a seven-day recovery period prior to shipping. Cores were wrapped in plastic bags and wet paper towels with a minimal amount of seawater, shipped in insulated coolers to the environmental simulation aquarium facility at San Francisco State University’s Estuary & Ocean Science Center in Tiburon, CA, and immediately placed in experimental aquaria. All corals were collected and exported under applicable permits from the National Park of American Samoa (NPSA-2006-SCI-0001) and the Department of Marine and Wildlife Resources and imported under the authority of the US Fish and Wildlife Service.

### Coral acclimation conditions

The cores from each source colony were divided into two groups of 15 and held in two separate experimental aquaria at a constant temperature of 28 ± 0.5 ºC and average irradiance of 260 µmol quanta m^-2^ sec^-1^ (12hr light/dark cycle) for a 28 day recovery period. Algal growth was removed from the nylon bolts daily during the recovery period using a toothbrush. Following the recovery period, corals were exposed to one of two different thermal conditions for 35 days: “variable” or “stable”. In the variable aquarium, temperatures fluctuated between 27 and 32 °C during the afternoon of each day (mean temperature = 28.2 °C) while the other aquarium was set to remain stable at 28.5 ºC (Fig 1). The specific temperatures and amplitude of the treatments were chosen to reflect average water temperature and daily range extremes of natural summer temperature profiles of the forereef and back reef sites (Barshis et al., 2010; Craig et al., 2001; Smith et al., 2008). Due to equipment malfunction, there were two days of the acclimation when the stable aquarium and variable aquarium reached the same high temperature and three days where the stable aquarium reached temperatures below that of the variable aquarium (Fig S1). After the 35-day acclimation period, coral growth, photophysiology, and protein stress biomarkers were assessed.

**Figure 1.**
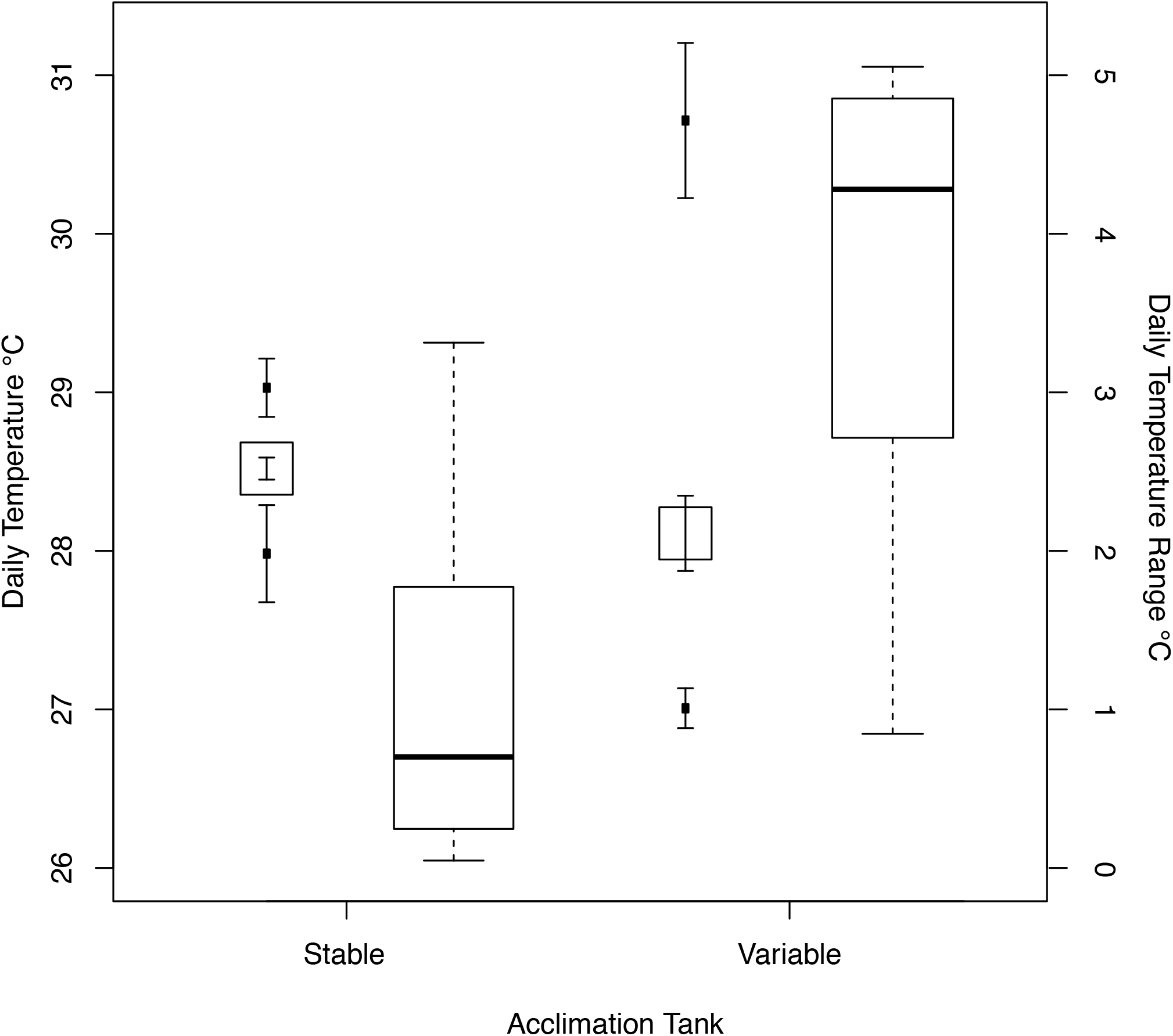
Daily mean (squares), minimum and maximum (dark circles) ± 95% confidence intervals of each acclimation aquarium during the 35-day acclimation period (left-hand axis) and boxplot of daily temperature range (right-hand axis) of each acclimation aquarium.

**Figure 2.**
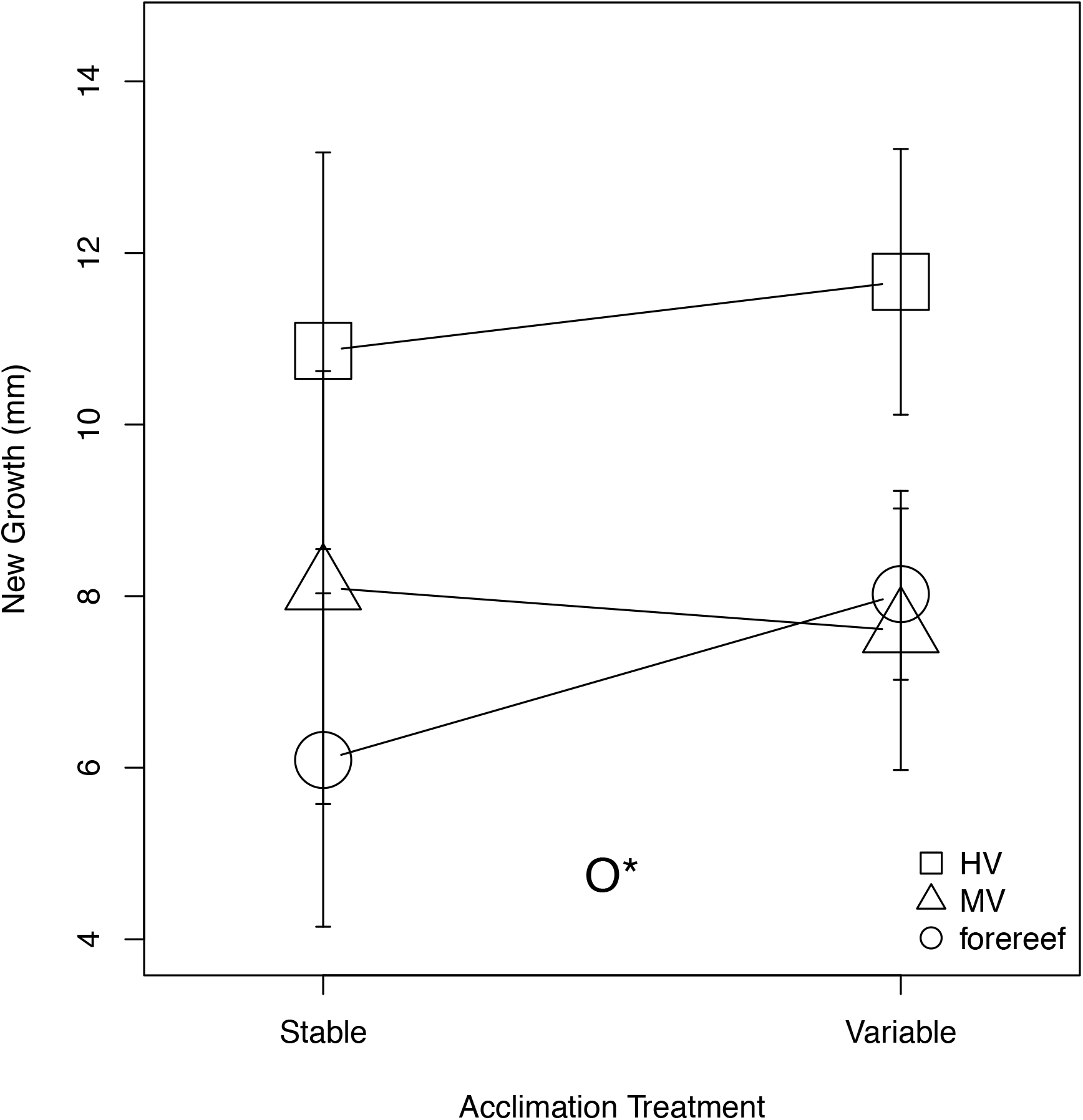
Linear tissue extension measurements taken after the 35-day acclimation period. Values are category means ± 1 SD. Statistical significance at p < 0.05 (*)is presented for comparison of source colony origins (O).

We acknowledge that it would have been preferable to directly replicate the acclimation tanks and treatments, however it was both logistically and financially prohibitive to do so. Each system cost many thousands of dollars to achieve such manipulable temperature control and the acclimation period (28 + 36 days) was of such an extended duration that successive field collections and trials were unable to be performed. Additionally, both aquaria had a constant flow of water from the same 5,000 L recirculating water source, likely preventing any substantial differences in water chemistry between tanks. Each tank was continuously fed from this water source throughout the experiment at a flow rate of ∼4 tank volumes per day. This same system has been used successfully in other published studies (e.g., Paganini et al., 2014). Furthermore, we believe the concordance between this study’s lab-based results and those of prior field experiments demonstrating strong fixed effects of origin in this species and minimal effects of acclimation treatment corroborate the assertion of little to no confounding influence of the single tank replicates on the results of the study.

### Growth

New tissue growth was measured as the distance the growth margin had extended down the sides of each coral core since original sampling; measured linearly down the four cardinal sides of each core using calipers. The four measurements were averaged and analyzed using a single average value for each individual core.

### Photophysiology

Chlorophyll *a* fluorescence of *Symbiodinium* sp. was measured using a pulse-amplitude modulated (PAM) fluorometer (DIVING-PAM, Walz GmbH, Germany). PAM fluorometry is a rapid, non-invasive technique which assesses the photosynthetic efficiency of photosystem II (PSII) reaction centers which can be used as a proxy for assessing the health of the symbiotic association (Fitt et al., 2001). DIVING-PAM parameters and measurements were made following a previous study (Piniak and Brown, 2009); initial fluorescence measurements (*F*) were between ∼150–400 units and maximum fluorescence (F’*m*) was measured using a saturating light pulse (0.8 s, ∼8000 umol quanta m^-2^ s^-1^). Maximum quantum yield [(*Fm* - *Fo*)/*Fm*, or *Fv / Fm*] was measured for dark-adapted samples at the end of each experimental day 45 min after all lights had been turned off.

### Thermal challenge

After the 35-day acclimation period, a “temperature ramp” was performed. This consisted of placing five replicate cores from all source colonies and treatments in the variable temperature aquarium (baseline 27, peak 32 ºC) for one day and subsequently raising the baseline and peak temperatures by 2 °C every 24 hrs for four additional days with a final temperature fluctuation of 35 – 40 ºC and total experimental duration of 120 hrs (Figs. S1, 3). PAM measurements were taken each day at 21:15 after 45 min of dark-adaptation. A single core from each source colony and acclimation treatment was sacrificed for protein analyses following PAM measurements each night, flash frozen in liquid nitrogen, and stored at –80 °C until analyzed as described below.

**Figure 3.**
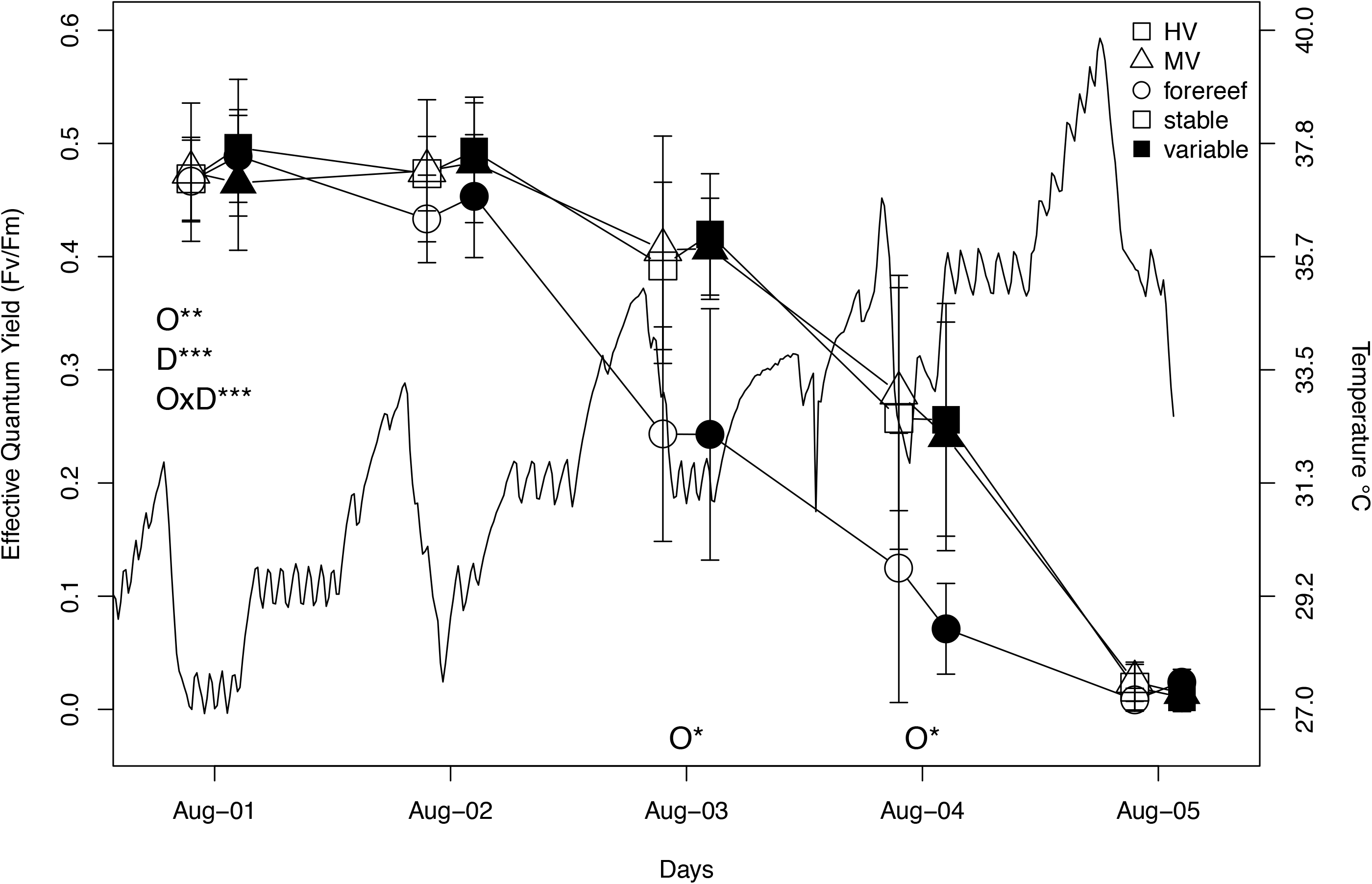
Pulse Amplitude Modulated fluorometry (PAM) measured maximum quantum yield (Fv/Fm) of corals during 5 days of the ramping temperature exposure. PAM measurements were taken at the end of the experimental day following ≥ 45 minutes of dark adaptation. The experimental temperature profile is shown as the solid black line and on the right-hand axis. Squares, triangles, and circles represent source colonies from the High Variability (HV) pool, Medium Variability (MV) pool, and forereef (FR) respectively and open and shaded symbols are for stable and variable acclimation treatments, respectively. Values are category means ± 1 SD. Statistical significance at p < 0.05 (*), p < 0.01 (**), and p < 0.001 (***) is presented for overall comparisons of source colony origin (O), acclimation treatment (A), day (D), and the various interactions (e.g., OxA) along the left hand y-axis, while within-day contrasts are presented along the x-axis.

### Protein biomarkers: Hsp70 and ubiquitin-conjugates

Each coral core was flash frozen in liquid nitrogen and the tissue layer (up to 1 cm below surface) was removed with bone cutting pliers and placed in a pre-frozen, 50 ml stainless steel mixing jar (Glennmills, Clifton, NJ). The tissue and skeleton of each tissue layer was crushed using a TissueLyser^®^ (Qiagen, Valencia, CA) at 25 rpm for 5 s, and the powdered samples were transferred to individual 2.5 ml cryovials and stored at –80 ºC until further analyses.

Between 280–380 mg of crushed tissue was added to a prechilled 2 ml microcentrifuge tube before adding 750 µL of chilled 50 mM phosphate buffer (K_2_HPO_4_ + KH_2_PO_4_; pH 7.8). Samples were vortexed and centrifuged at 2,000 x *g* for 5 min to separate out host and algal endosymbiont (*Symbiodinium*) tissue fractions. The supernatant (host fraction) was removed and placed on ice while the remaining pellet (skeletal debris and *Symbiodinium* fraction) was washed three times with fresh phosphate buffer before re-suspension in a final volume of 500 μL of phosphate buffer, sonicated for 5 min, and briefly centrifuged to remove skeletal debris. Aliquots were removed from both host and *Symbiodinium* fractions and stored at –80 ºC until further analyses.

Levels of heat shock protein 70 (hsp70) and ubiquitin-conjugated proteins were assessed via western blot for both host and *Symbiodinium* protein fractions as described previously (Barshis et al., 2010). All samples were assayed in triplicate and a single average concentration per sample was analyzed.

### Host genetic analyses

To assess the potential influence of host genotype on physiological responses, the internal transcribed spacer region (ITS) of nuclear ribosomal DNA was amplified and sequenced from each individual source colony. Primer sequences, polymerase chain reaction conditions, and sequencing methods were performed as described previously (Barshis et al., 2010). Resulting sequences were inspected using Sequencher version 4.5 (Gene Codes Corp., Ann Arbor, MI) and aligned using Bio-edit (Hall, 2001) and by eye. Population genetic structure was estimated using an analysis of molecular variance (AMOVA) in Arlequin 2.0 (Schneider, 2000). A molecular phylogenetic network was constructed using the median-joining algorithm and maximum parsimony post-processing calculation in NETWORK ver 4.5.0.0 (Fluxus Technology Ltd., Polzin and Daneschmand, 2003).

### Statistical analyses

Within a common garden framework, comparisons between transplant groups are designed to assess acclimation potential versus genetic/epigenetic control over the response variables. Comparisons between acclimation treatments examine environmental effects (i.e., phenotypic plasticity), while comparisons between source colony origins and individuals examine potential genetic or epigenetic influence on the response variables (DeWitt and Scheiner, 2004; Schluter, 2000; Smith et al., 2007).

Growth, photosynthetic efficiency, and western blot biomarker levels were assessed from field collections prior to shipping (field baseline), following the acclimation to the differing temperature profiles of the two experimental aquaria (acclimation baseline), and during the temperature ramp. For the field baseline and acclimation baseline tests, all variables were tested against the fixed factors of source colony origin and acclimation treatment in a two-way ANOVA (aov) with source colony individual (i.e., genotype) included as a random factor. Post-hoc analyses of significant main effects were computed using the lsmeans function in R v3.2.2 (R_Core_Team, 2015). Individual clonal replicates within time points were averaged prior to the ANOVA and plotting to avoid pseudoreplication. Assumptions of normality and homoscedasticity were tested via the shapiro.test and fligner.test functions in R v3.2.2 (R_Core_Team, 2015), respectively. For comparisons across time points, a repeated measures framework was used incorporating source colony identity (i.e., individual genotype) as a unit of repeated measure, allowing for a between-subjects test of origin and within-subjects tests of acclimation and day. Post-hoc analyses of multiple comparisons were computed using the lsmeans function in R v3.2.2 (R_Core_Team, 2015).

## Results

### Initial acclimation: temperature

The stable treatment had a slightly higher mean temperature and lower standard deviation than the variable treatment (28.54 ± 0.63 and 28.15 ± 1.20 ºC for the stable and variable tanks respectively; Figs. 1, S1). On average, the daily range of the variable tank was 11.84 times greater than the daily range of the stable tank. Of the 33 acclimation days for which temperature records were available, the variable tank had a daily range greater than 3 ºC on 24 days (73 %), while the daily range of the stable tank exceeded 3 ºC on only one day due to a heater malfunction (Fig. S1). Irradiance levels did not appear to differ between the two tanks with an average irradiance of 263 and 259 μmol quanta m^-2^ sec^-1^ for the stable and variable tank respectively.

### Initial acclimation: growth

New tissue extension during the acclimation period was affected by source colony origin (p = 0.0106; Fig. 2, Table S1). HV source colonies grew fastest overall, with an average tissue extension of 10.85 mm ± 2.72 SD and 11.66 mm ± 2.23 SD in the stable and variable tanks respectively compared to MV source colonies (8.07 mm ± 2.62 SD and 7.60 mm ± 2.01 SD) and forereef source colonies (6.06 mm ± 2.00 SD and 8.01 mm ± 1.80 SD) for the stable and variable treatments respectively. There was no significant difference in growth between acclimation treatments for corals from any origin (p = 0.0977; Fig. 2, Table S1).

### Temperature ramp: photophysiology

Measurements of maximum quantum yield (*Fv/Fm*) were significantly affected by source colony origin, day, and an origin x day interaction (p = 0.0036, <0.0001, <0.0001 respectively; Fig. 3, Table S2). On days three and four of the temperature ramp, corals from the thermally stable forereef had markedly lower effective quantum yield (Fv/Fm) compared to back-reef corals regardless of acclimation treatment (Fig. 3, Table S2). There was no effect of acclimation treatment throughout the temperature ramp. By the end of the ramp (day 5), corals from all populations had little to no fluorescence signature (Figs. 3, Table S2).

### Hsp70 and ubiquitin-conjugates: *Symbiodinium* fraction

Hsp70 levels in the *Symbiodinium* fraction of field-collected samples were different among origins (p = 0.0259, Table S3A), with forereef levels 3.5 times lower than MV corals (p = 0.0249; Fig. 4, Table S3A). Ubiquitin-conjugate levels were also different among origins (p = 0.0352; Fig. 5, Table S4A), with forereef levels 10.3 times lower than HV corals (p = 0.0439; Fig. 5, Table S4A). Following acclimation, both origin, acclimation, and origin * acclimation effects were observed (p = 0.0398, p < 0.0001, and p = 0.0093 respectively; Table S3B) in *Symbiodinium* hsp70 levels, with 3.3 times lower levels in the stable vs variable acclimation treatment (p < 0.0001; Table S3B) and 10.2 times higher in the HV vs. MV or forereef variable acclimated corals (p = 0.0029, p = 0.0032 for MV and forereef contrasts respectively; Table S3B). *Symbiodinium* ubiquitin-conjugates were not different amongst origins or acclimation treatments following acclimation (Fig. 5, Table S4B). During the temperature ramp, a mix of origin and acclimation effects were observed, with variable and contrasting responses across groups throughout the experiment (Figs. 4, 5, Tables S3C, S4C).

**Figure 4.**
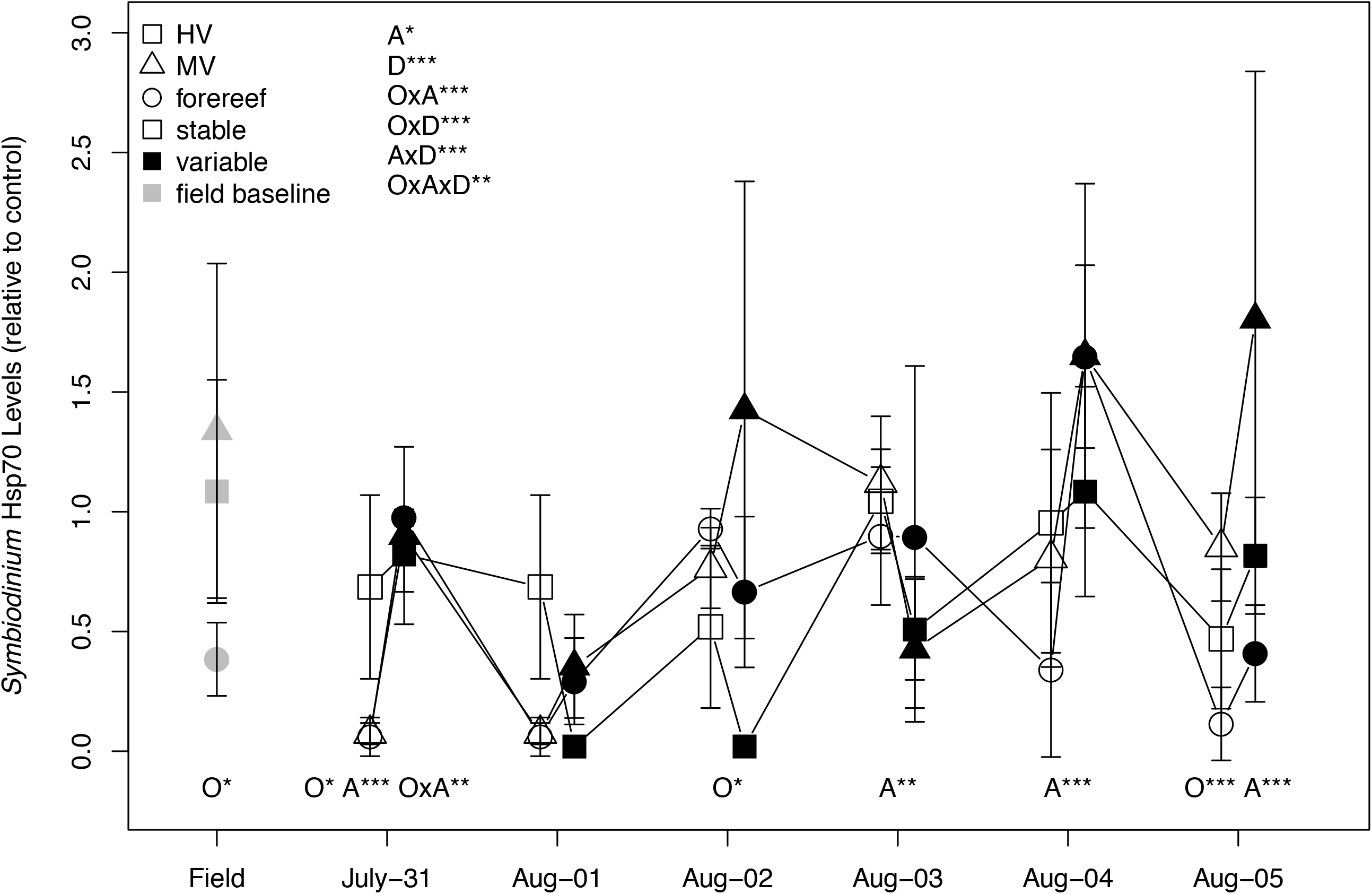
*Symbiodinium* heat shock protein 70 levels across the entire sampling period: field baseline, post-acclimation, and days 1–5 of the temperature ramp. All values are relative to a single control extract. Values are category means ± 1 SD. Symbols and significance values are denoted as in Figure 3.

**Figure 5.**
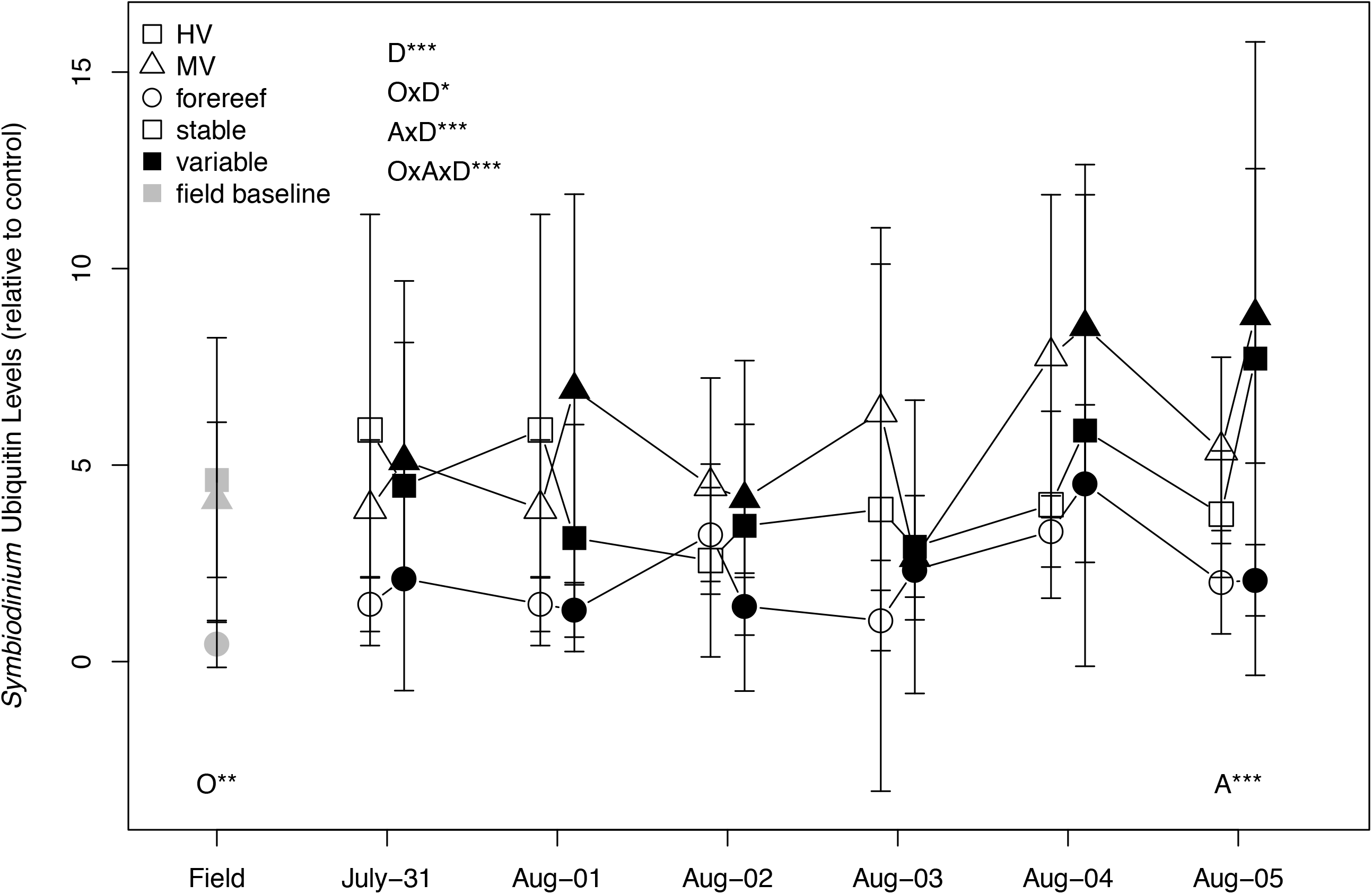
*Symbiodinium* ubiquitin-conjugate protein levels across the entire sampling period: field baseline, post-acclimation, and days 1–5 of the temperature ramp. All values are relative to a single control extract. Values are category means ± 1 SD. Symbols and significance values are denoted as in Figure 3.

### Hsp70 and ubiquitin-conjugates: Host fraction

Neither host hsp70 nor ubiquitin-conjugate protein levels were different in the field-collected samples (Figs. 6, 7, Tables S5A, S6A). Following acclimation, host hsp70 levels were similar amongst origins but 1.4 times higher on average in stable-acclimated corals (p = 0.0029; Fig 5, Table S5B), while ubiquitin-conjugates were 2.2 times lower on average in stable-acclimated corals (p = 0.0012; Fig 7, Table S6B). Host hsp70 levels were 2.9 and 2.5 times higher in HV vs. forereef corals on days 2 and 4 of the heat ramp respectively (p = 0.0129, p = 0.0404; Fig 6, Table S5C), and there was a significant origin x day interaction in host ubiquitin-conjugate levels (p = 0.0080), though no significant individual contrasts (Fig 7, Table S6C).

**Figure 6.**
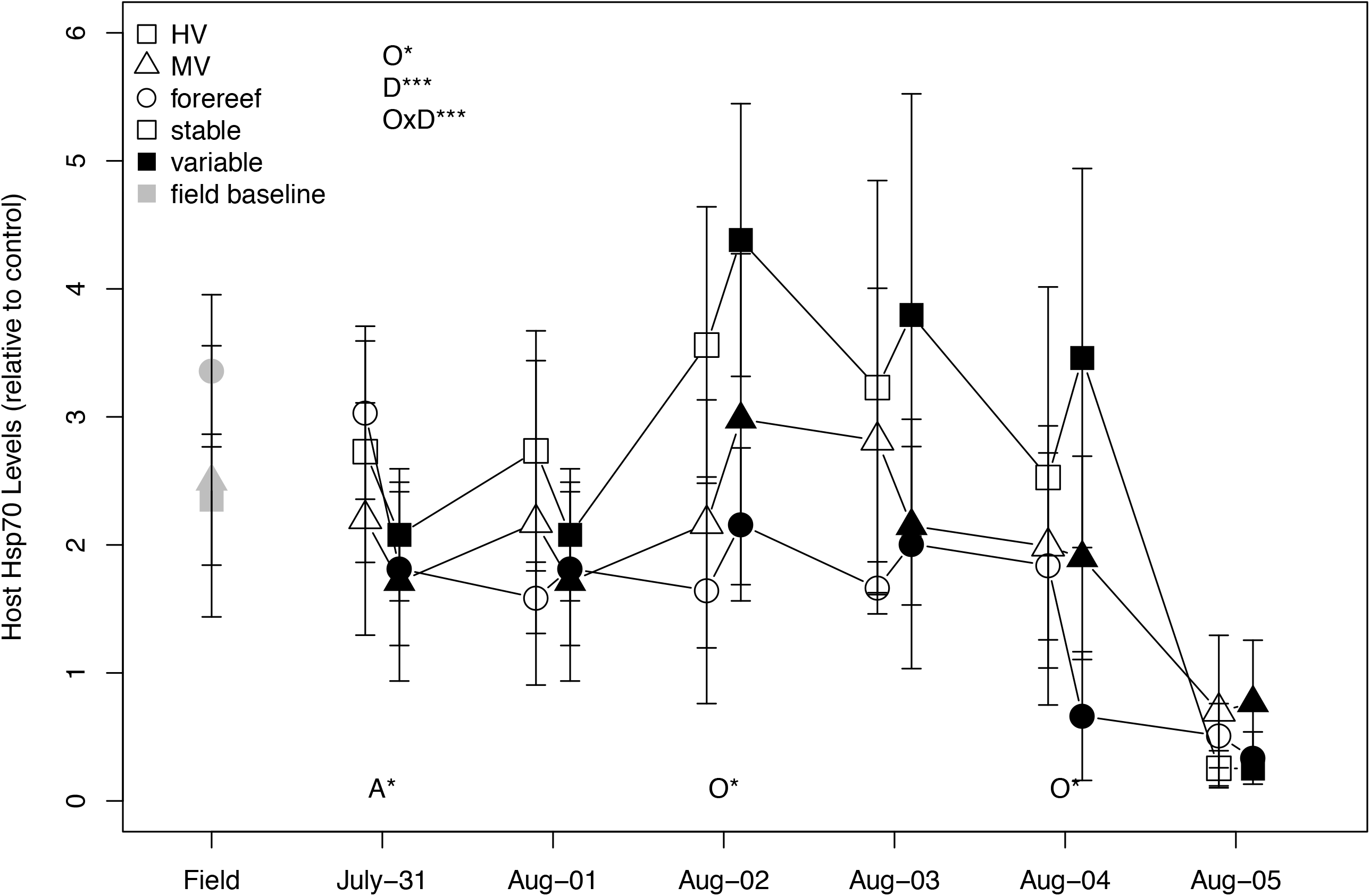
Coral host heat shock protein 70 levels across the entire sampling period: field baseline, post-acclimation, and days 1–5 of the temperature ramp. All values are relative to a single control extract. Values are category means ± 1 SD. Symbols and significance values are denoted as in Figure 3.

**Figure 7.**
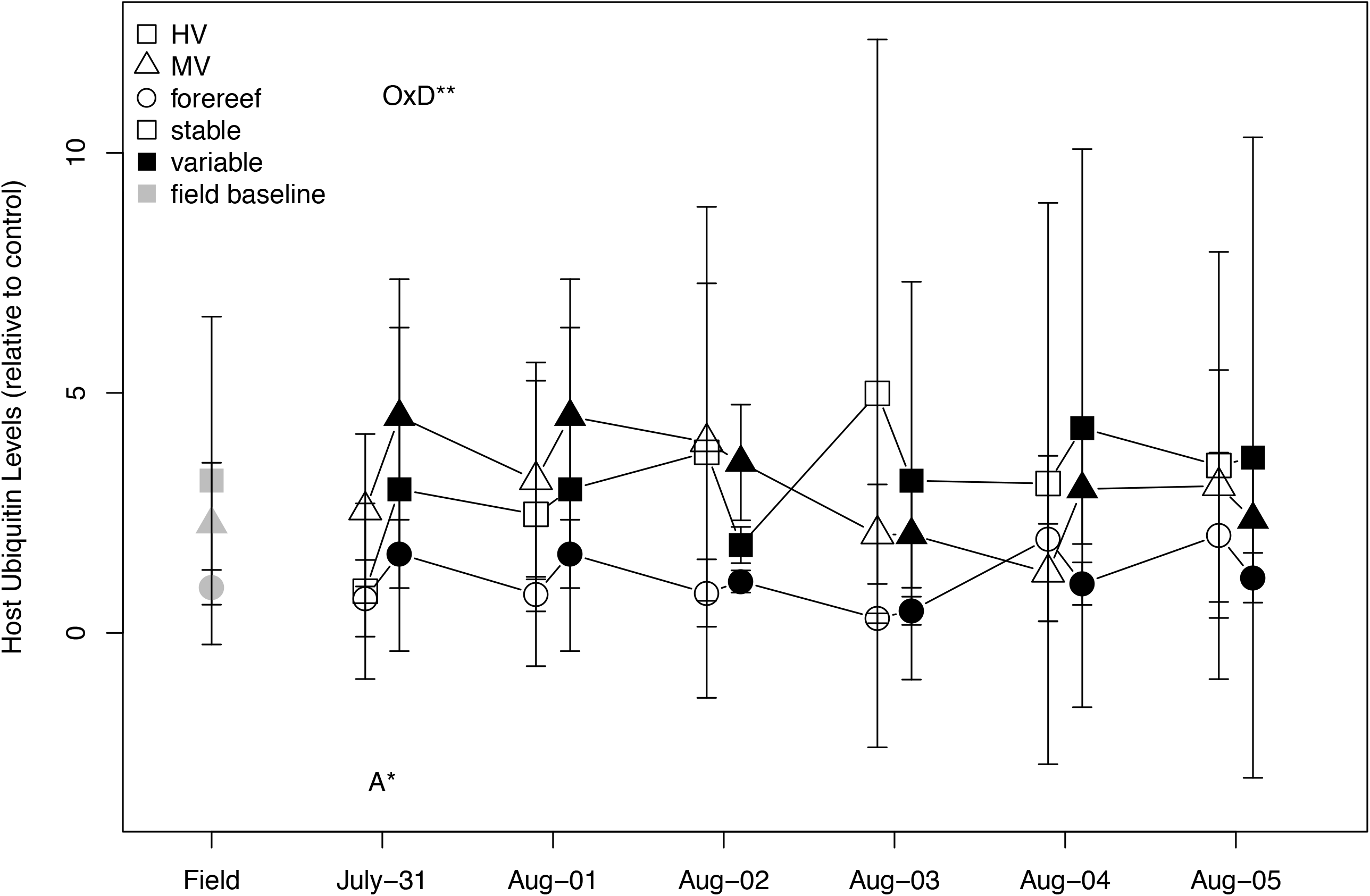
Coral ubiquitin-conjugate protein levels across the entire sampling period: field baseline, post-acclimation, and days 1–5 of the temperature ramp. All values are relative to a single control extract. Values are category means ± 1 SD. Symbols and significance values are denoted as in Figure 3.

### Host genetic analyses

A 368 base pair (bp) fragment of the internal transcribed spacer region (ITS) was amplified from 15 individuals (n=5 per origin) and subsequently cloned for a total of 77 cloned sequences (NCBI accession numbers xxxxxx – xxxxxx). These 77 sequences were comprised of 28 unique haplotypes: one shared between the HV and MV pools, one shared between the HV pool and forereef, and eight, nine, and nine unique to the HV, MV, and forereef sites respectively (Fig. 8). An analysis of molecular variance (AMOVA) revealed significant population subdivision among all three populations (*F_ST_* = 0.2061, p < 0.0001). Pairwise *F_ST_* comparisons were highest between the MV pool and the other two populations (*F_ST_* = 0.2483 and 0.2319, p < 0.0001 for HV and forereef respectively), while the HV pool and forereef showed lower but still significant subdivision (*F_ST_* = 0.0509, p < 0.03). This was qualitatively evident in the phylogenetic network construction, which showed a more explicit separation between MV haplotypes versus HV and forereef haplotypes (Fig. 8).

**Figure 8.**
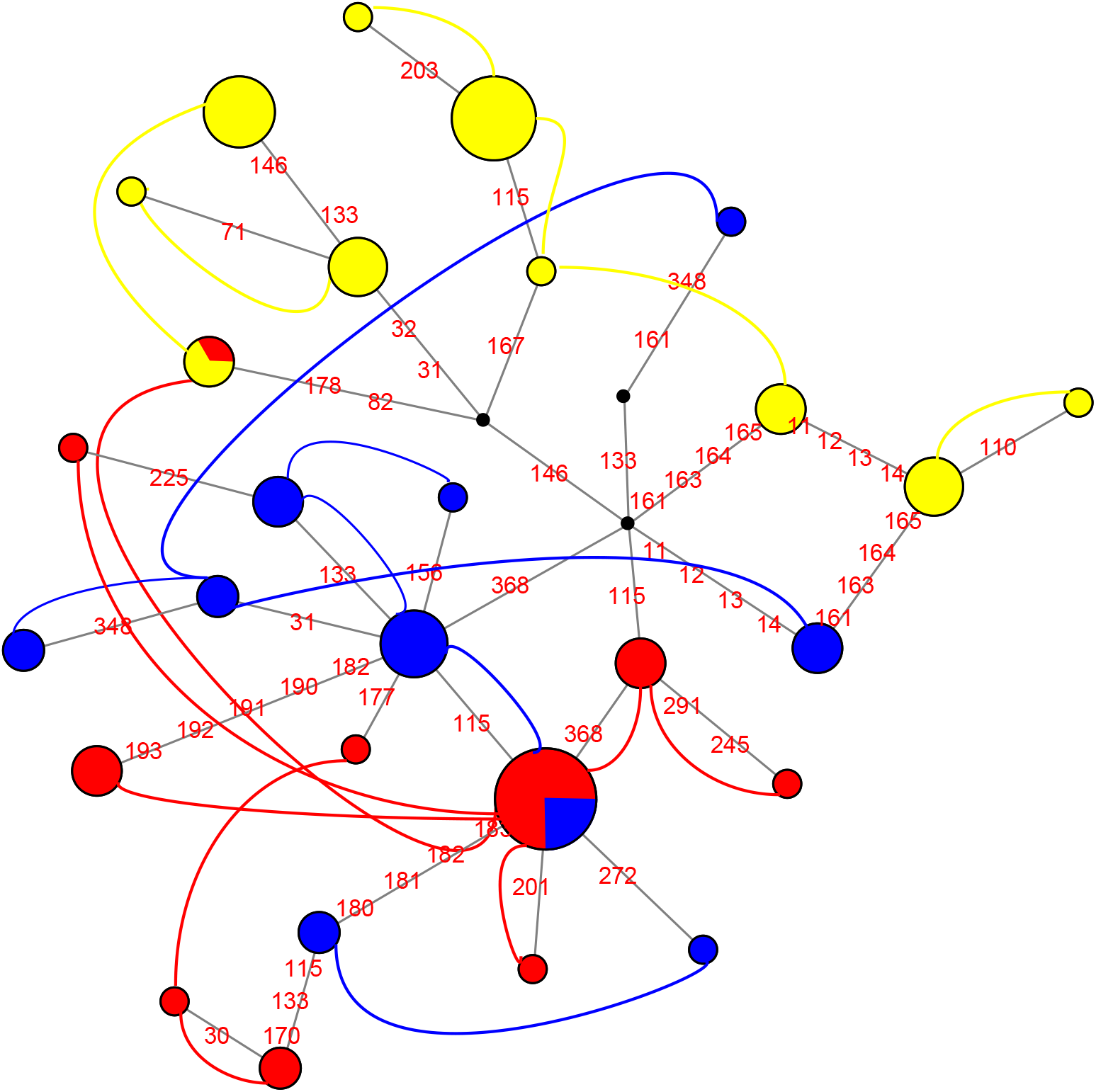
Maximum parsimony phylogenetic network reconstruction of ITS rDNA haplotypes drawn in NETWORK ver 4.5.0.0 (Fluxus Technology Ltd.; Polzin & Daneschmand 2003). Haplotypes shown in red, yellow, and blue are from HV, MV, and forereef populations respectively. Diameter of circles at each node is proportional to the number of individuals with identical sequences. Haplotypes that co-occur in the same individual are connected by colored curves. Mutations are shown in red on each branch with the number corresponding to the base pair position, hypothetical intermediate haplotypes are designated by black circles. NCBI accession numbers for all sequences used in this study are XXXXX-XXXXX.

## Discussion

### The influence of high-frequency variability on coral physiological tolerance limits

Here, we found that increasing the amount of high-frequency thermal variability (i.e., diurnal or shorter time-scales) for 36 days of acclimation had little to no effect on coral growth, photophysiology, thermal tolerance, or protein biomarker response (Figs. 2–7). The predominant signal in our data was that of source population origin, in that corals from back reef habitats (HV and MV) with consistent high-frequency variability in thermal and other environmental characteristics grew faster and had elevated thermal tolerance limits compared to corals from the more thermally stable forereef, regardless of acclimation treatment. Taken together, these results suggest real differences in thermal tolerance limits between back reef corals that have routinely been exposed to high-frequency environmental variability and forereef corals native to a less-variable environment. The disparity between the lack of acclimation effects and strong origin effects speaks to the potential for chronic exposure to high-frequency variability to exert differential selection pressure over very small spatial scales (< 5 km).

The most widely-used models of coral bleaching impacts and thermal tolerance differences rely on island-scale or regional level data (e.g., the 5 km pixel width of NOAA Coral Reef Watch; Heron et al., 2016). However, our findings demonstrate substantial differences in coral thermal tolerances across hundreds of meters to a few kilometers. This follows previous results from Ofu corals in the highest variability back reef habitats showing meter-scale differences in increased prevalence of heat-tolerant clade D *Symbiodinium* (e.g., Acropora spp., Pocillopora spp., Pavona sp.; Cunning et al., 2015; Oliver and Palumbi, 2009), constitutive turning-on of genes involved in cellular stress defense (Barshis et al., 2013), acclimation gains in thermal tolerance following 12+ months of exposure to the HV pool (Palumbi et al., 2014), and small-scale (< 5 km) genetic differentiation of coral hosts consistent with local adaptation (Barshis et al., 2010; Bay and Palumbi, 2014).

A number of other studies across the globe have found similar small-scale differences in physiological tolerance limits between corals from habitats with contrasting amounts of short-term environmental variability. For example, *Porites astreoides* corals from inshore environments with high-frequency thermal variability in the Florida Keys bleached less during thermal stress (Kenkel et al., 2013), demonstrated increased flexibility in gene expression modulation (Kenkel and Matz, 2016), and increased growth rates that were heritable between generations (Kenkel et al., 2015) compared to corals from lower variability offshore sites (∼ 7 km away). Similarly, Pineda et al. (2013) found decreased mortality in *Stylophora pistillata* on protected (shoreward) vs. exposed (seaward) sides of reefs in the central Red Sea following a natural bleaching event in 2010. Despite being separated by < 300 m, the protected sides of the reefs had greater high-frequency thermal variability than exposed sites presumably due to decreased wind-driven mixing (Pineda et al., 2013). Similar increased stress tolerance was observed in inshore vs. offshore populations of *Montastrea annularis* in Belize (Castillo and Helmuth, 2005), which was subsequently linked to long-term declines in growth rates in offshore populations of this species over the past few decades (Castillo et al., 2012). A recent large-scale meta-analysis of in-situ temperature records and bleaching surveys from 5 reef regions around the globe found that greater amounts of high-frequency temperature variability were correlated with reduced bleaching severity and bleaching prevalence (Safaie et al., 2018), suggesting the trends observed in the various single-site, single-species studies may be valid at the global and whole-reef community scales.

There are a few notable exceptions to this pattern, however, with high variability and low variability populations of *Acropora palmata* and *Porites astreoides* in the Cayman Islands exhibiting a nearly identical response to increased heat and pCO_2_ exposure (Camp et al., 2016), and exposure to greater high-frequency thermal variability eliciting bleaching rather than resilience in *Pocillopora meandrina* and *Porites rus* in Moorea, French Polynesia (Putnam and Edmunds, 2011). While the specific threshold above which high-frequency variability increases resilience and/or the tipping points between beneficial exposures versus chronic stress remain to be determined, our data corroborate a growing body of evidence from multiple ocean basins, coral species, genera, and habitat types suggesting a mostly beneficial role of high-frequency variability in increasing coral resilience to thermal stress. Thus, it is conceivable that that differing degrees of environmental variability may exert divergent selection pressures across these small-scales and drive adaptive differentiation.

### Is temperature variability really the most important driver?

Despite the overall effects of source colony origin, however, we found little evidence that acclimation to high-frequency temperature variability altered thermal tolerance limits in this species. In contrast to the lack of acclimation observed herein, multiple studies of *Acropora* spp. have demonstrated increased thermal tolerance following short-term (days to weeks) exposure to elevated temperatures. *Acropora nana* from a single back-reef population on Ofu exposed to variable temperatures (29–33 °C) bleached less and had a muted gene expression response compared to corals acclimated to 29 °C after just 7–11 days of exposure to the variable thermal regime (Bay and Palumbi, 2015). Similarly, *Acropora millepora* preconditioned to a 10-day mild stress (3 °C below the experimentally determined bleaching threshold) bleached less during subsequent heat stress than non-preconditioned corals (Bellantuono et al., 2012b) and exhibited a muted gene expression response as well (Bellantuono et al., 2012a), similar to that seen in variable acclimated *Acropora nana* (Bay and Palumbi, 2015) and HV *A. hyacinthus* (Barshis et al., 2013) in Ofu. Lastly, *Acropora aspera* preconditioned to a 48 hr pre-stress (31 °C) bleached less and maintained elevated photosynthetic efficiency during a subsequent 6-day heat stress (34 °C) compared to non-preconditioned corals (Middlebrook et al., 2008).

Most prior thermal-acclimation work in corals has focused on branching species in the genus *Acropora*, due to their ubiquity on the reef and known variation in thermal sensitivity (e.g., Loya et al., 2001; van Woesik et al., 2011). In contrast, massive coral species such as *Porites lobata*, are thought to be more thermally tolerant due to greater tissue thicknesses (Loya et al., 2001), increased mass transfer rates (Loya et al., 2001; Nakamura and Van Woesik, 2001), and elevated metabolism (Gates and Edmunds, 1999) compared to most branching coral species (primarily *Acropora* and *Pocillopora*). Thus, as a species with a massive morphology, *Porites lobata* may have a greater innate temperature tolerance range to begin with, simply tolerating the environment when faced with new conditions versus the physiological acclimation seen in Acroporids. However, the consistent origin effects on growth, thermal tolerance, and cellular response suggests that the differing amounts of high-frequency variability in environmental characteristics between the back reef and forereef habitats do influence thermal tolerance limits in *P. lobata*, though perhaps over longer timescales than those under investigation.

Significant origin effects in common garden experiments are generally attributed to potential genotypic (i.e., adaptive) influence on the response variable (DeWitt and Scheiner, 2004; Sanford and Kelly, 2011; Schluter, 2000). However, long-term acclimatization, developmental plasticity, and/or epigenetics can similarly cause apparent origin effects. Corals are long-lived organisms, and based on the size (>1 m diameter) of the colonies used in this study, we roughly estimate the minimum age of the source colonies to be > 60 years old (based on > 500 mm radius and ∼8 mm/year growth rate sensu Houck et al., 1977; Potts et al., 1985). Decadal-scale ‘environmental memory’ was recently observed in the massive coral *Coelastrea aspera*, with former west sides of colonies (experimentally turned to face east) that had been previously exposed to high-irradiance levels retaining four times the *Symbiodinium* during a natural bleaching event compared to un-manipulated east-facing/low-irradiance sides of colonies; despite 10 years of conditioning to the low-irradiance eastern orientation and identical *Symbiodinium* phylotypes (Brown et al., 2015). This certainly raises the possibility that long-term conditioning to the high-frequency environmental variability of the Ofu back-reef could have long-lasting acclimation effects on *P. lobata* thermal tolerance limits that may not have been altered by our 36-day exposure.

However, we did observe acclimation effects on host and *Symbiodinium* protein biomarkers, particularly hsp70 (Figs. 4–7; Tables S3-S6). While differences across acclimation treatments were variable in magnitude and direction depending on the marker and day, the host hsp70 response demonstrated an interesting pattern relative to the fluorescence response. On the final day of the acclimation treatment, host hsp70 levels were lower in the variable vs. stable acclimated corals (Fig. 6, Table S5B), suggesting reduced need for chaperone activity following variable thermal exposure. However, the initial acclimation effect was supplanted by a strong origin effect with the greatest host hsp70 increase in HV corals on days 2 and 4 of the temperature ramp (Fig. 5, Table S5C), corresponding to the greater maintenance of photosynthetic efficiency in HV corals on days 3 and 4 (Fig. 3, Table S2). It is notable that a similarly rapid and higher induction of hsp70 was observed in back reef vs. forereef corals in our previous field study following transplantation (Fig. 4A from Barshis et al., 2010). Thus, it is tempting to speculate that the larger and more rapid hsp70 increases in HV corals during the temperature ramp might signify a higher capacity for maintenance of homeostasis under thermal stress. While not conclusive evidence for or against a mechanism of long-term acclimatization vs. local adaptation, the acclimation and origin effects in protein response observed herein and previously (Barshis et al., 2010) demonstrate the ability of these corals to respond to high-frequency thermal variability over short time-scales as well as potential evolutionary constraints on that ability related to population of origin.

Alternatively, the increased thermal tolerance limits of back-reef corals may have been influenced very early on either via developmental canalization post-settlement in the back-reef, parental effects, and/or epigenetic acclimatization. Both maternal effects and signatures of differential epigenetic modification have been recently observed in *Pocillopora damicornis*, with larvae from parents exposed to high temperature and *p* CO_2_ exhibiting metabolic acclimation during subsequent stress compared to larvae from un-exposed parents (Putnam and Gates, 2015) and increased levels of DNA methylation in adults following high *p* CO_2_ exposure (Putnam et al., 2016), suggesting that the observed larval acclimation could have been caused by epigenetic modification. In Ofu, however, larvae from back reef parents would have to settle/disperse back to the pool of origin for epigenetic modification from parents to positively affect the response of the offspring. If there was epigenetic modification of larvae from back reef parents but the larvae all dispersed outside the HV and MV pools, then there would be no positive contribution to the phenotype of the next generation.

While long-term acclimatization, parental effects, and/or epigenetic modification could explain the thermal tolerance differences between our populations, none of these processes would likely cause the genetic differentiation among populations seen here. The significant genetic subdivision among all three populations suggests the presence of a physical or environmental barrier to gene flow between the HV, MV, and forereef populations, strong divergent selection pressures, or potential cryptic species/genepools across the habitats in Ofu. Reduced connectivity across such a small spatial scale (∼500 m –1 km between HV and MV, and ∼5 km between HV/MV and the forereef) is unlikely to be due to a physical barrier alone, as the water masses in the back reef appear to be well-mixed during the daily high tide cycle and well-within the spatial range of dispersing larvae. Bay and Palumbi (2014) observed a similar pattern of genetic differentiation between HV and MV *Acropora hyacinthus*, though only at a subset of outlier single nucleotide polymorphisms (SNPs) putatively responding to selection. They posited a mechanism involving strong spatial balancing selection, wherein the contrasting environmental pressures of each habitat exert high selection pressure on settling coral larvae from a common gene pool (sensu a protected polymorphism via an environment x genotype association; Bay and Palumbi, 2014; Levene, 1953; Ravigné et al., 2004; Sanford and Kelly, 2011). van Oppen et al. (2018) found a similar pattern of differentiation and outlier loci separating reef flat and reef slope *Pocillopora damicornis* on Heron Island in Australia and posited a similar mechanism of environmentally driven selection. The ITS locus sequenced herein is unlikely to be a direct target of selection, though differentiation at this locus could be correlated with the specific gene targets responding to selection.

### Conclusions

The limited acclimation response, enhanced thermal tolerance capacity of back reef corals, differential biomarker response, and significant genetic differentiation observed in the present study are all consistent with a model of post-settlement selection and adaptation of coral genotoypes to the greater amount of high-frequency environmental variability in the MV and HV pools. However, the lack of acclimation in thermal tolerance limits following 35 days of exposure to *temperature* variability alone, calls into question whether differences in high-frequency temperature exposures among habitats are the driving force behind these differences. Differences in the amount of high-frequency temperature variability remains the common factor across the multiple experiments on Ofu (Barshis et al., 2013; Barshis et al., 2010; Bay and Palumbi, 2014; Craig et al., 2001; Cunning et al., 2015; Oliver and Palumbi, 2011; Palumbi et al., 2014; Smith et al., 2007), the Red Sea (Pineda et al., 2013), Florida Keys (Kenkel et al., 2013; Kenkel et al., 2015; Kenkel and Matz, 2016), meso-american barrier reef (Castillo and Helmuth, 2005; Castillo et al., 2012), Australia (van Oppen et al., 2018), and the variety of sites examined in Safaie et al. (2018). Future research should focus on assessing the potential influences of other environmental drivers on the observed differences in thermal limits, as well as the relative contributions of long-term acclimatization and/or developmental canalization. Additionally, the magnitude of *F_ST_* differentiation observed herein makes it difficult to contextualize the scale of genetic differentiation across populations. Future in-depth genetic analysis of massive *Porites* populations from a variety of habitat types may provide a clearer picture of the potential for cryptic genotype x environment associations in this taxonomic group.

## Acknowledgements

Thanks to the many field and laboratory assistants that made this research possible; to M. Mizobe, L. Smith, T. Waterson, C. Tomas-Miranda, the Stillman Lab at SFSU, P. Craig and the staff at the National Park of American Samoa especially the late Fale Tuilagi, and Vaoto Lodge for field and lab support; Z. Forsman for primer sequences, advice, and assistance; and to anonymous reviewers for valuable comments. This is contribution number #### of the Hawai’i Institute of Marine Biology. The use of trade, firm, or corporation names in this publication does not constitute an official endorsement or approval of any product or service to the exclusion of others that may be suitable.

## Data Accessibility

All raw data, analyses, and scripts are available as an electronic notebook: (https://github.com/BarshisLab/Poriteslobata-thermal-adaptation). DNA sequences have also been deposited in NCBI Genbank (accession #’s XXXXXXX).

## Author Contributions

DJB, JHS, CB, RJT, and RDG conceived and designed the research. DJB and JHS conducted the research. CB, RJT, RDG, and JHS funded the research. DJB and JHS performed the analyses. DJB, JHS, CB, RJT, and RDG wrote the paper.

## Supplementary Figure Legends

**Figure S1.** Experimental tank temperatures measured every 15 min during the 36-day acclimation period. The variable tank is shown in red and the stable tank in blue.

